# Evaluation of gene-based family-based methods to detect novel genes associated with familial late onset Alzheimer disease

**DOI:** 10.1101/242545

**Authors:** Maria Victoria Fernández, John Budde, Jorge Del-Aguila, Laura Ibañez, Yuetiva Deming, Oscar Harari, Joanne Norton, John C Morris, Alison Goate, NIA-LOAD family study group, NCRAD, Carlos Cruchaga

## Abstract

Gene-based tests to study the combined effect of rare variants towards a particular phenotype have been widely developed for case-control studies, but their evolution and adaptation for family-based studies, especially for complex incomplete families, has been slower. In this study, we have performed a practical examination of all the latest gene-based methods available for family-based study designs using both simulated and real datasets. We have examined the performance of several collapsing, variance-component and transmission disequilibrium tests across eight different software and twenty-two models utilizing a cohort of 285 families (N=1,235) with late-onset Alzheimer disease (LOAD). After a thorough examination of each of these tests, we propose a methodological approach to identify, with high confidence, genes associated with the studied phenotype with high confidence and we provide recommendations to select the best software and model for family-based gene-based analyses. Additionally, in our dataset, we identified *PTK2B*, a GWAS candidate gene for sporadic AD, along with six novel genes (*CHRD, CLCN2, HDLBP, CPAMD8, NLRP9, MAS1L*) as candidates genes for familial LOAD.

## 1 Introduction

Alzheimer disease (AD) is a complex condition for which almost 50% of its phenotypic variability is due to genetic causes; yet, only 30% of the genetic variability is explained by known markers (Ridge et al. 2016). GWAS studies have identified more than 20 risk loci (Lambert et al. 2013); and sequencing studies have identified additional genes harboring low frequency variants with large effect size (*TREM2, PDL3, UNC5C, SORL1, ABCA7*, (Sims et al. 2017)). Recent studies also indicate that Late-Onset AD (LOAD) families are enriched for genetic risk factors (Cruchaga et al. 2017). Therefore studying those families may lead to the identification of novel variants and genes (Cruchaga et al. 2014; Guerreiro et al. 2013).

Current consensus is that the missing heritability for complex traits and AD may be hidden under the effect of rare variants with low to moderate effect on disease risk (Frazer et al. 2009; Manolio et al. 2009; Cirulli and Goldstein 2010). The rarity of these markers requires specific study designs and statistical analysis for their detection. The simplest approach to detect rare variants for association is to test each variant individually using standard contingency table and regression methods. But due to the few observations of the rare minor allele at a specific variant, the statistical power to detect association with any rare variant is limited; hence, extremely large samples are required and a more stringent multiple-test correction applies as compared to common variants (Bansal et al. 2010; B. Li and Leal 2008). It has been acknowledged that the best alternative is to collapse sets of pre-defined candidate rare variants within significant units, usually genes (gene-based sets) (Lee et al. 2014; Neale and Sham 2004). Collapsing tests work under the framework of giving each variant a certain weight and perform summation of weights through all variants within the region; depending on the weights and how summation is performed there are four major types of gene-based methods: collapsing tests, variance-component tests, and combined tests (Lee et al. 2014). Collapsing tests, analyze whether the overall burden of rare variants is significantly different in cases compared to controls by regressing disease status on minor allele counts (MAC). The Cohort Allelic Sum Test (CAST) is a dominant genetic model that assumes that the presence of any rare variant increases disease risk (Morgenthaler and Thilly 2007); whereas the Combined Multivariate and Collapsing (CMC) method, collapses rare variants in different MAF categories and evaluates the joint effect of common and rare variants through Hoteling’s test (Li and Leal 2008). However, neither CAST nor CMC tests allow correcting for directional effect. The Variable Threshold (VT) test instead allows for both trait-increasing and trait-decreasing variants; it selects optimal frequency thresholds for burden tests of rare variants and estimates p-values analytically or by permutation (Price et al. 2010). Variance-componence methods test for association by evaluating the distribution of genetic effects for a group of variants while appropriately weighting the contribution of each variant. The sequence kernel association test (SKAT) casts the problem in mixed models (Lee et al. 2014), and in the absence of covariates, SKAT reduces to C-alpha test. (Neale et al. 2011). Finally, collapsing and variance component tests can be combined into one statistical method, the SKAT-O approach (Lee et al. 2012), which is statistically efficient regardless of the direction and effect of the variants studied.

All these methods were initially designed for unrelated case-control study designs; but given the rarity of these variants, large datasets are required to achieve statistical power. (Laird and Lange 2006). Alternatively, family-based studies in which several family members share the same phenotype may provide more statistical power than regular case-controls studies (Li et al. 2006; Cirulli and Goldstein 2010; Ott et al. 2011; Kazma and Bailey 2011). Pioneering methods were designed for testing nuclear families, trios or sibships (Ionita-Laza et al. 2013; Horvath et al. 2001; Laird et al. 2000; De et al. 2013; Ott et al. 2011). However, considering the late-onset nature of Alzheimer disease it is often difficult to obtain genetic information from parents (to conform trios), or nuclear family units. The usual pedigree in familial LOAD corresponds to incomplete, large familial units (**Figure 1**). Most of the initial software for gene-based family-based studies were not suitable for complex pedigrees like those observed in Alzheimer studies, but in recent years a plethora of methods have been developed that take into account complex family structure in gene-based calculations. Among the software that take into account large pedigrees we find SKAT (Wu et al. 2011), FSKAT (Yan et al. 2015), GSKAT (Wang et al. 2013), RV-GDT (Chen et al. 2009), EPACTS (http://genome.sph.umich.edu/wiki/EPACTS), FarVAT (Choi et al. 2014), PedGene (Schaid et al. 2013) and RareIBD (Sul et al. 2016).

**Figure 1.**
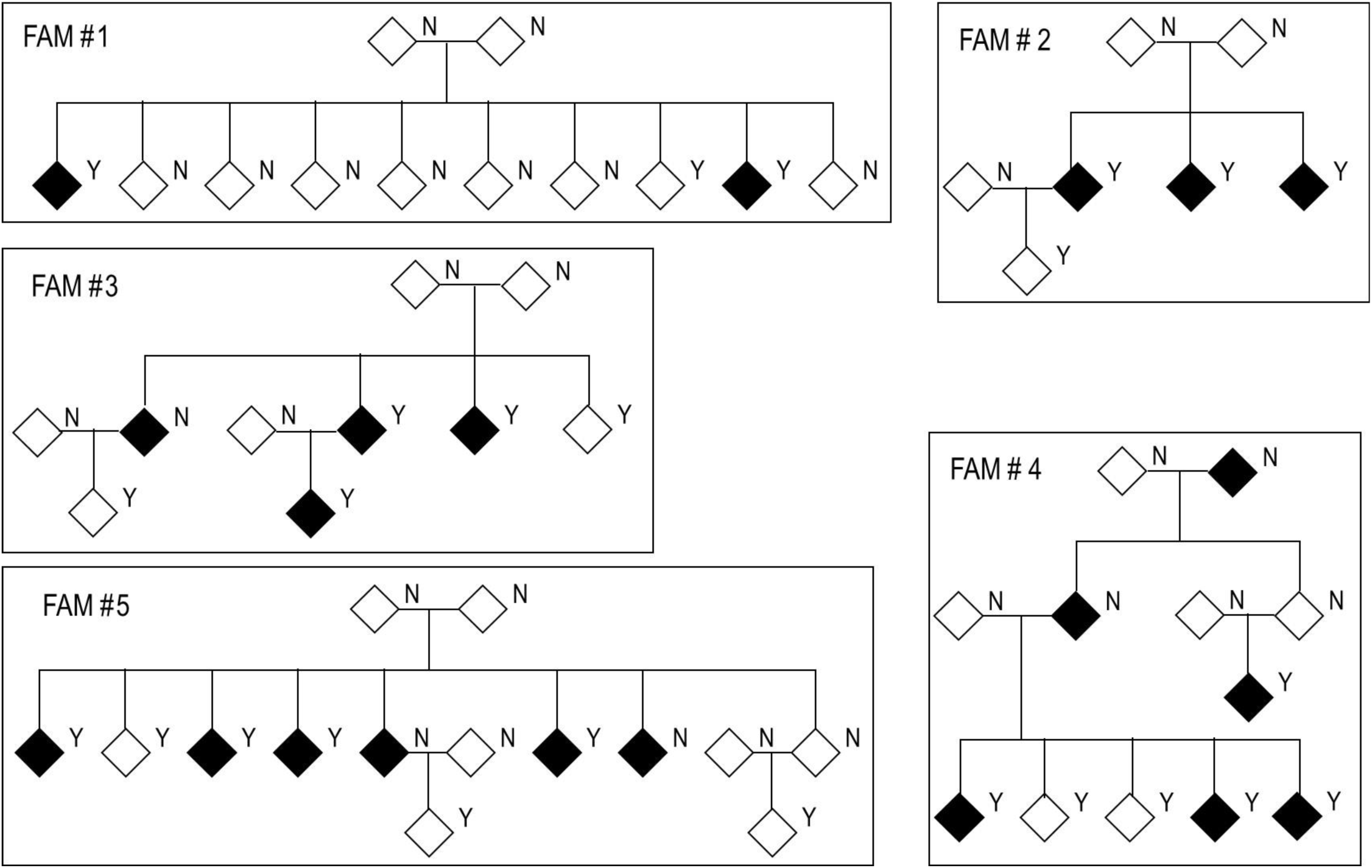
Structure of families used in this study. Black diamonds represent cases and white diamonds represent controls. Y: genetic data available. N: no genetic data available.

In this study, we wanted to evaluate the performance of the eight most common gene-based family-based methods available using a real dataset, over 250 multiplex families affected with Alzheimer disease, under different conditions and models. We simulated multiple scenarios in which a candidate variant perfectly segregates with disease status to rank the different programs and models. We also tested the performance of these tests at evaluating known causal genes for AD in our cohort. Finally, we performed genome-wide analysis to evaluate the power of each of these tests. Altogether, we discuss the pros and cons of each method that can be very informative for other investigators performing similar analyses: complex diseases in complex, incomplete, large families. We want to emphasize that although this work is centered on AD, the information extracted from this work can be equally applied to other complex traits. Finally, based on the results from the methods analyzed, we present some candidate genes for AD.

## 2 Materials and Methods

### 2.1 Cohort

The LOAD families included in this study originated from two cohorts: Washington University School of Medicine (WUSM) cohort and ADSP cohort.

#### 2.1.1 WUSM cohort

Samples from the Washington University School of Medicine (WUSM) cohort were recruited by either the Charles F. and Joanne Knight Alzheimer's Disease Research Center (Knight ADRC) at the WUSM in Saint Louis or the National Institute on Aging Genetics Initiative for Late-Onset Alzheimer’s Disease (NIA-LOAD). This study was approved by each recruiting center Institutional Review Board. Research was carried out in accordance with the approved protocol. Written informed consent was obtained from participants and their family members by the Clinical and Genetics Core of the Knight ADRC. The approval number for the Knight ADRC Genetics Core family studies is 201104178. The NIA-LOAD Family Study has recruited multiplex families with two or more siblings affected with LOAD across the United States. A description of these samples has been reported previously (Wijsman et al. 2011) (Fernández et al. 2017; Cruchaga et al. 2012). We selected individuals for sequencing from families in which APOEε4 did not segregate with disease status, and in which the proband of the family did not carry any known mutation in *APP, PSEN1, PSEN2, MAPT, GRN* or *C9orf72* (described previously (Cruchaga et al. 2012)).

#### 2.1.2 ADSP cohort

The Alzheimer's Disease Sequencing Project (ADSP) is a collaborative work of five independent groups across the USA that aims to identify new genomic variants contributing to increased risk for LOAD. (https://www.niagads.org/adsp/content/home). During the discovery phase, they generated whole genome sequence (WGS) data from members of multiplex LOAD families, and whole exome sequence (WES) data from a large case-control cohort. These data are available to qualified researchers through the database of Genotypes and Phenotypes (https://www.ncbi.nlm.nih.gov/gap Study Accession: phs000572.v7.p4).

The familial cohort of the ADSP consists of 582 individuals from 111 multiplex AD families from European-American, Caribbean Hispanic, and Dutch ancestry (details about the samples are available at NIAGADS). We downloaded raw data (.sra format) from dbGAP for 143 IDs (113 cases and 23 controls) from 37 multiplex families of European-American ancestry that were incorporated with the WUSM cohort.

### 2.2 Sequencing

Samples were sequenced using either whole-genome sequencing (WGS, 12%) or whole-exome sequencing (WES, 88%). Exome libraries were prepared using Agilent’s SureSelect Human All Exon kits V3 and V5 or Roche VCRome (Table 2). Both, WES and WGS samples were sequenced on a HiSeq2000 with paired ends reads, with a mean depth of coverage of 50× to 150× for WES and 30× for WGS. Alignment was conducted against GRCh37.p13 genome reference. Variant calling was performed separately for WES and WGS following GATK’s 3.6 Best Practices (https://software.broadinstitute.org/gatk/best-practices/) and restricted to Agilent’s V5 kit plus a 100bp of padding added to each capture target end. We used BCFTOOLS (https://samtools.github.io/bcftools/bcftools.html) to decompose multiallelic variants into biallelic prior variant quality control. Variant Quality Score Recalibration (VQSR) was performed separately for WES and WGS, and for SNPs and INDELs. Only those SNPs and indels that fell within the above 99.9 confidence threshold, as indicated by WQSR, were considered for analysis; variants within low complexity regions were removed from both WES and WGS and variants with a depth (DP) larger than the average DP + 5 SD in the WGS dataset were removed. At this point SNPs and indels from WES and WGS datasets were merged into one file. Non-polymorphic variants and those outside the expected ratio of allele balance for heterozygosity calls (ABHet=0.3-0.7) were removed. Additional hard filters implemented included quality depth (QD ≥7 for indels and QD≥2 for SNPs), mapping quality (MQ≥40), fisher strand balance (FS≥200 for indels and FS≥60 for SNPs), Strand Odds Ratio (SOR≥10 for Indels and SOR≥3 for SNPs), Inbreeding Coefficient (IC ≥-0.8 for indels) and Rank Sum Test for relative positioning of reference versus alternative alleles within reads (RPRS≥-20 for Indels and RPRS≥-8 for SNPs) (**Figure S1**). We used PLINK1.9 (https://www.cog-genomics.org/plink2/ibd) to remove variants out of Hardy Weinberg equilibrium (p-value <1×10^-6^), with a genotype calling rate below 95%, with differential missingness between cases vs controls, WES vs WGS, or among different sequencing platforms (p-value<1×10^-6^).

Samples with more than 10% of missing variants (four samples) and whose genotype data indicated a sex discordant from the clinical database (three samples) were removed from dataset. Individual and familial relatedness was confirmed using identity-by-descent (IBD) calculations, an existing GWAS dataset for these individuals, and the pedigree information. Because many of the ADSP families were also recruited from the NIA-LOAD repository there is a certain overlap (48 individuals) between the WUSM and the ADSP familial cohorts; we kept the duplicated pair that had better genotyping rate after QC. Principal Component Analysis (PCA) was calculated to corroborate ancestry and restrict our analysis to only samples from European American origin. Functional impact and population frequencies of variants were annotated with SnpEff (Cingolani et al. 2012). For this analysis, only SNVs with a minor allele frequency (MAF) below 1%, as registered in ExAC (Lek et al. 2016),were taken into account.

We excluded families carrying a known pathogenic mutation in any of the Mendelian genes for Alzheimer disease, Frontotemporal Dementia, or Parkinson disease (Fernández et al. 2017). We restricted the selection of families to those families with at least one case and one control in the family, and we excluded any participants initially diagnosed as AD but that turned into other after pathological examination. Finally, our dataset consisted of 1235 non-hispanic whites (NHW), 824 cases and 411 controls, from 285 different families (Table 1, Table S1).

**Table 1.** Demographic data for the familial dataset employed in this study.

|  | N | *Age ±<br>SD | *Age<br>range | % Fe | %<br>APOE4 |
| --- | --- | --- | --- | --- | --- |
| Cases | 824 | 73 ±7 | 48-99 | 63% | 73% |
| Controls | 411 | 83 ± 9 | 39-104 | 59% | 51% |
| Total | 1235 | 77 ± 10 | 39-104 | 61% | 65% |

\* Age At Onset (AAO) for cases and Age at Last Assessment (ALA) for controls.

**Table 2.** Number of samples for which whole genome sequencing (WGS) or whole exome sequencing (WES) was performed, with detail of the exon library kits employed in this study.

| Exon library kit | WGS | WES |
| --- | --- | --- |
| WGS | 153 |  |
| Agilent's SureSelect Human All Exon kits V3 | 0 | 28 |
| Agilent's SureSelect Human All Exon kits V5 | 0 | 665 |
| Roche VCRome | 0 | 389 |
| Total | 153 | 1082 |

### 2.3 Study design & analysis.

The goal of this study was to test the performance and power of different gene-based family-based methods available to date, using a real dataset consisting of 1,235 non-hispanic white individuals from 285 families densely affected with AD. We set up three different scenarios to test (**Figure 2**). First, using the real phenotype and pedigree structure of 25 from the 285 families, we generated a synthetic dataset with multiple variants and families with perfect segregation. Second, we evaluated different variant-combinations for the *APOE* gene. Third, we performed genome-wide gene-based analysis accounting only for non-synonymous SNPs with a MAF < 1%. For each one of these scenarios we evaluated the performance of the different gene-based methods (collapsing, variance-component, and transmission disequilibrium) from the following family-based packages: SKAT (Wu et al. 2011), FSKAT (Yan et al. 2015), GSKAT (Wang et al. 2013), RVGDT (He et al. 2017), EPACTS (http://genome.sph.umich.edu/wiki/EPACTS), FarVAT (Choi et al. 2014), PedGene (Schaid et al. 2013), RareIBD (Sul et al. 2016). Some of these software offer the option to run different gene-based algorithms; e.g. GSKAT, EPACTS, FarVAT or PedGene can run collapsing and variance-component tests; therefore, we ran a total of 25 models (**Table 3**). The details of each one of these scenarios are described next.

**Figure 2.**
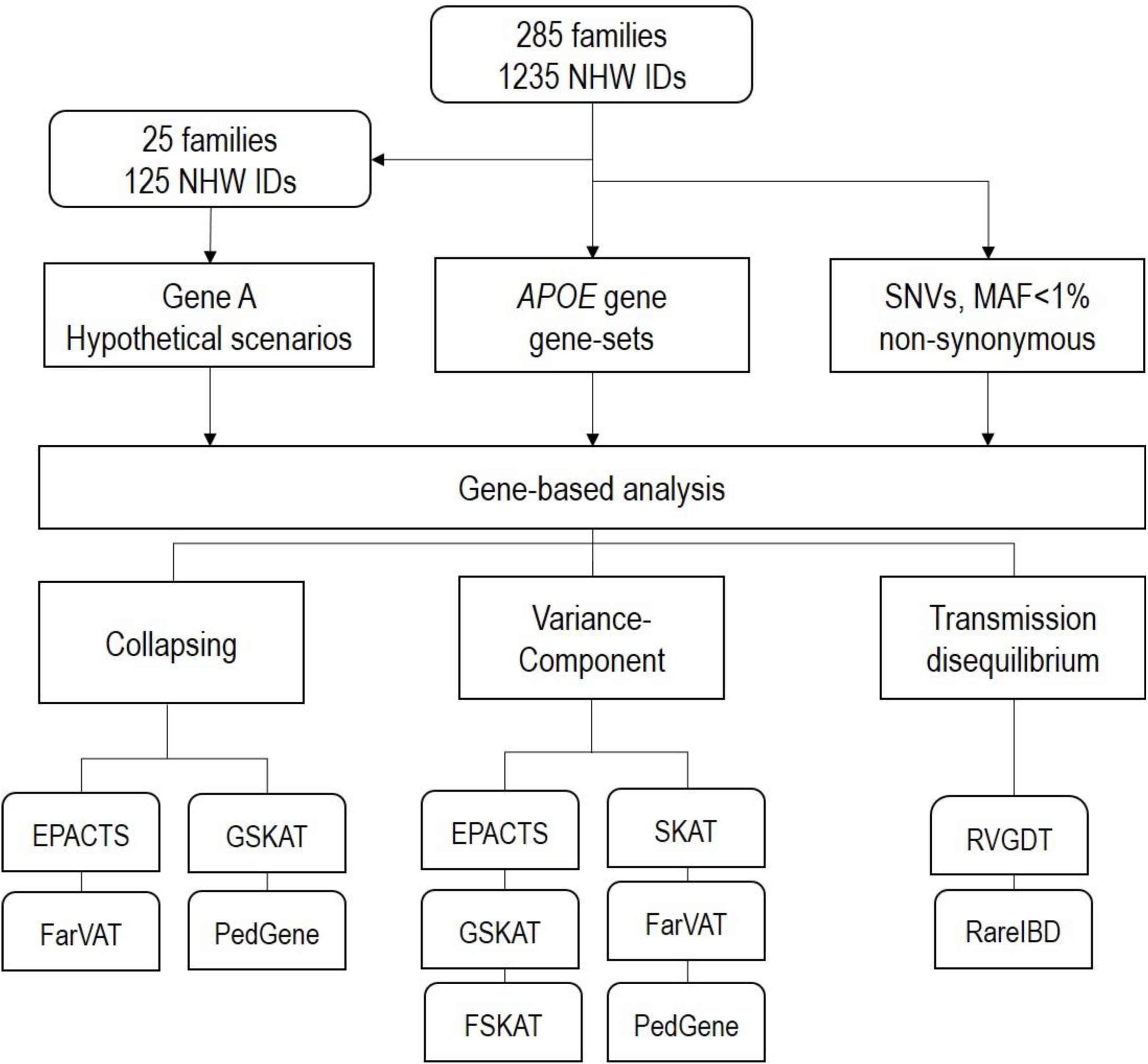
Schematic design of the analysis performed in this study.

**Table 3.** Relationship of programs and models tested according to their main features and kinship matrix that they use.

|  | Collapsing |  |  | Variance-component |  | Combined | Transmission-disequilibrium | Kinship |  |  |
| --- | --- | --- | --- | --- | --- | --- | --- | --- | --- | --- |
|  | Burden | CMC | VT | C-ALPHA | SKAT | SKATO |  | BN | IBS | Ped |
| EPACTS |  | X | X |  | X |  |  | X |  |  |
| RVGDT |  |  |  |  |  |  | X |  |  |  |
| SKAT-v2 |  |  |  |  | X |  |  | X | X | X |
| GSKAT | X |  |  |  | X |  |  |  |  | X |
| FSKAT |  |  |  |  | X |  |  |  |  | X |
| FarVat-Adj | X | X |  | X |  | X |  |  |  |  |
| FarVat-BLUP | X | X |  | X |  | X |  |  |  |  |
| Pedgne | X |  |  |  | X |  |  |  |  |  |
| RareIbd |  |  |  |  |  |  | X |  |  |  |

#### 2.3.1 Simulated data

We selected 25 representative families from our entire dataset for which there was genotypic data for three to seven members (Table S2). We used the existing family structure and phenotype of these families, and a simulated gene called “GENE-A” containing five variants. We generated several scenarios in which different numbers of families presented perfect segregation with disease status for a variant in GENE-A (Table 4 and Table S2). First, we considered a scenario in which only the first five families of the dataset were included in the analyses, and each family presented a different perfectly segregating variant of GENE-A (scenario 5 family carriers (FC) and 0 non-carriers (FNC): 5FC×0FNC). Second, we generated additional scenarios in which we kept the same five families carrier of segregating variants in GENE-A, and added five (scenario 5FC×5FNC), ten (scenario 5FC×10FNC), 15 (scenario 5FC×15FNC), and 20 (scenario 5FC×20FNC) families that were not carriers of any variant in GENE-A. Then, we considered four scenarios of 25 families in which each new scenario added families who were carriers of a segregating variant in GENE-A. We started with the scenario 5FC×20FNC, then we simulated ten families carriers and 15 families non-carriers (scenario 10FC×15FNC), 15 families carries and 10 families non-carriers (scenario 15FC×10FNC), 20 families carriers and five families non-carriers (scenario 20FC×5FNC) and concluded with a scenario in which all 25 families were carriers of one, of the possible five, segregating variant in GENE-A (scenario 25FC×0FNC). We tested each one of these scenarios with all previously mentioned gene-based methods and software to evaluate their power to associate perfect segregating variants with disease.

**Table 4.** Representation of the segregation pattern of the simulated gene. One (1) means that all cases within the family are carriers of the variant. Zero (0) means that the variant is not present in that family.

|  | GENE-A |  |  |  |  |
| --- | --- | --- | --- | --- | --- |
|  | SNP1 | SNP2 | SNP3 | SNP4 | SNP5 |
| Fam1 | 1 | 0 | 0 | 0 | 0 |
| Fam2 | 0 | 1 | 0 | 0 | 0 |
| Fam3 | 0 | 0 | 1 | 0 | 0 |
| Fam4 | 0 | 0 | 0 | 1 | 0 |
| Fam5 | 0 | 0 | 0 | 0 | 1 |
| Fam6 | 1 | 0 | 0 | 0 | 0 |
| Fam7 | 0 | 1 | 0 | 0 | 0 |
| Fam8 | 0 | 0 | 1 | 0 | 0 |
| Fam9 | 0 | 0 | 0 | 1 | 0 |
| Fam10 | 0 | 0 | 0 | 0 | 1 |
| Fam11 | 1 | 0 | 0 | 0 | 0 |
| Fam12 | 0 | 1 | 0 | 0 | 0 |
| Fam13 | 0 | 0 | 1 | 0 | 0 |
| Fam14 | 0 | 0 | 0 | 1 | 0 |
| Fam15 | 0 | 0 | 0 | 0 | 1 |
| Fam16 | 1 | 0 | 0 | 0 | 0 |
| Fam17 | 0 | 1 | 0 | 0 | 0 |
| Fam18 | 0 | 0 | 1 | 0 | 0 |
| Fam19 | 0 | 0 | 0 | 1 | 0 |
| Fam20 | 0 | 0 | 0 | 0 | 1 |
| Fam21 | 1 | 0 | 0 | 0 | 0 |
| Fam22 | 0 | 1 | 0 | 0 | 0 |
| Fam23 | 0 | 0 | 1 | 0 | 0 |
| Fam24 | 0 | 0 | 0 | 1 | 0 |
| Fam25 | 0 | 0 | 0 | 0 | 1 |

**Table 4.**
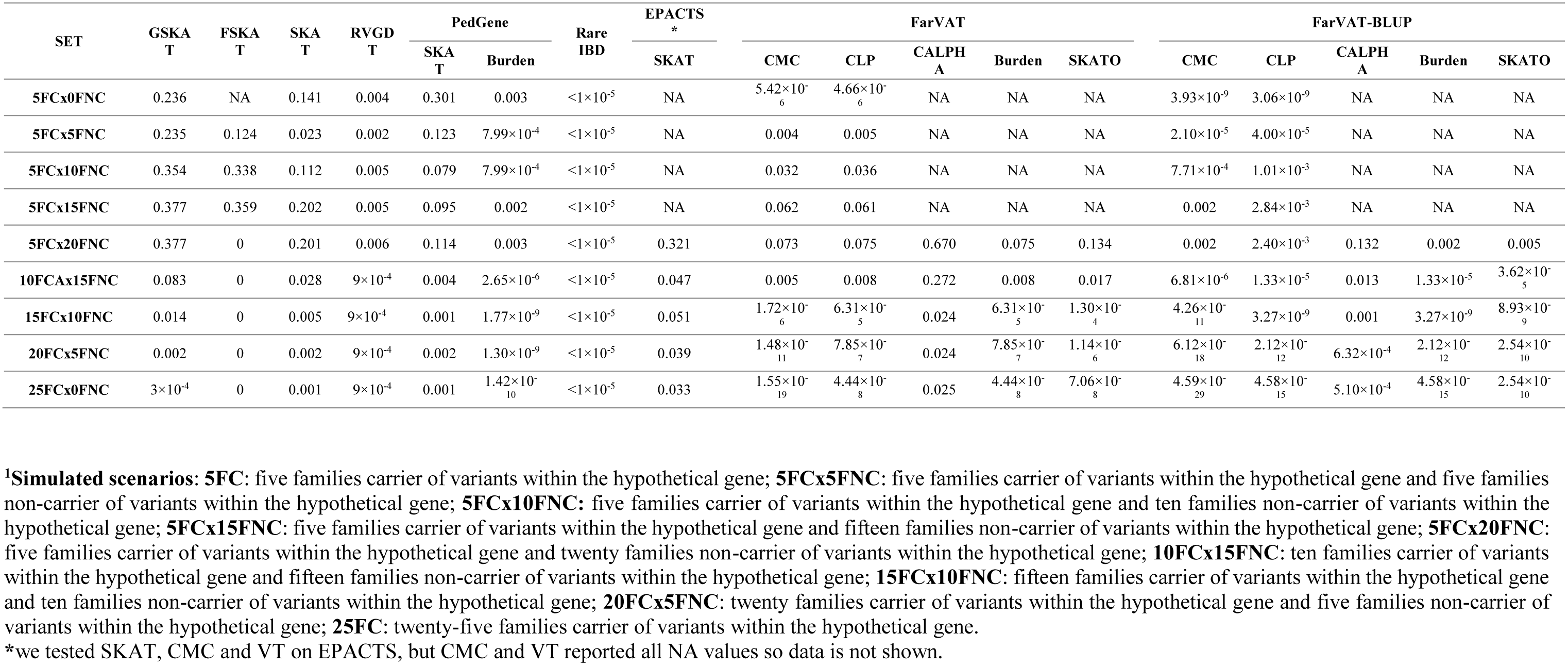
Gene-based p-values for the simulated dataset under different scenarios for the gene-based methods tested in the subset of 25 families.

#### 2.3.2 Candidate genes

*APOE* is the largest genetic risk factor for Alzheimer’s disease. The allelic combination of two SNPs, rs429358 (APOE 4; 19:45411941:T:C) and rs7412 (APOE 2: 19:45412079:C:T), determines one of the three major isoforms of APOE protein, ε2, ε3 or ε4. The dosage of these isoforms determines a person’s risk to suffer AD, from having a protective effect APOE ε2/ε2 (OR 0.6) or ε2/ε3 (OR 0.6) to different degrees of increased risk according to the number of copies of the ε4 allele (ε2/ε4, OR 2.6; ε3/ε4, OR 3.2; ε4/ε4, OR 14.9) (Farrer et al. 1997). We tested the power of all previously mentioned gene-based methods and software to detect association of *APOE* gene with disease in our entire dataset (N=1,235) under different conditions. We first tested all polymorphic variants (nonsynonymous with MAF <1%) in the *APOE* gene, second we tested only those variants considered to have a high or moderate effect on the protein including rs429358 and rs7412, and then we tested high and moderate variants alone, and finally tested rs429358 and rs7412 alone.

#### 2.3.3 Genome-wide analyses

We performed gene-based burden analysis on a genome-wide level in our entire dataset (families n=285; samples N=1,235) to evaluate the power of each of the previously mentioned methods to detect novel genes significantly associated with disease; only single nucleotide variants (SNVs) with a minor allele frequency equal or below 1%, based on the EXAC dataset (Lek et al. 2016) (MAF ≤ 1%), and with a predicted high or moderate effect, according to SnpEff (Cingolani et al. 2012) were included in the analysis. Quantile-Quantile (QQ) plots from gene-based p-values were generated with the R package “ggplot2” (Wickham 2009). We also evaluated the correlation between these methods using Pearson correlation (Pc) and Spearman correlation (Sc) tests on the log of the p-value using R v3.4.0 (R Core Team 2017). Pc evaluates the linear relationship between two continuous variables whereas Sc evaluates the monotonic relationship between two continuous or ordinal variables.

### 2.4 Software tested

A companying supporting file (**Supplementary material**) provides a summary of the code employed to run each of the programs described below.

#### 2.4.1 GSKAT

GSKAT (Wang et al. 2013) is among the first R packages to come out with the goal of extending burden and kernel-based gene set association tests for population data to related samples with binary phenotypes. To handle the correlated or clustered structure in the family data, GSKAT fits a marginal model with generalized estimated equations (GEE). The basic idea of GEE is to replace the covariance matrix in a generalized linear mix model (GLMM) with a working covariance matrix that reflects the cluster dependencies. Accordingly, GSKAT blends the strengths of kernel machine methods and generalized estimating equations (GEE), to test for the association between a phenotype and multiple variants in a SNP set. We ran GSKAT correcting for sex and first two PCs.

#### 2.4.2 SKAT

The sequence kernel association test SKAT (Wu et al. 2011) is an R package initially designed for case-control analysis. Later they incorporated the Efficient Mixed-Model Association eXpedited (EMMAX) algorithm (Zhou and Stephens 2012; Kang et al. 2010) that allows for performing family-based analysis. EMMAX simultaneously corrects for both population stratification and relatedness in an association study by using a linear mixed model with an empirically estimated relatedness matrix to model the correlation between phenotypes of sample subjects. The efficient application of EMMAX algorithm depends on appropriate estimate of the variance parameters. Relatedness matrices can be calculated based on pedigree structure or estimated from genotype data. For the latter, different methods have been proposed. Relatedness can be estimated using those alleles that have descended from a single ancestral allele, i.e. those that are Identical by Descent (IBD), or using the Balding-Nichols (BN) method (Balding and Nichols 1995) which explicitly models current day populations via their divergence from an ancestral population specified by Wright's *F_st_* statistic. We ran SKAT v1.2.1, on R v3.3.3, using option SKAT_Null_EMMAX correcting for sex and first two PCs and we tested four different kinship matrices: pedigree, IBS, BN and a BN based kinship matrix (HR) that EPACTS software constructs (**Table S3**).

#### 2.4.3 FSKAT

FSKAT (Yan et al. 2015), also an R package, is based on a kernel machine regression and can be viewed as an extension of the sequence kernel association test (SKAT and famSKAT) for application to family data with dichotomous traits. FSKAT is based on a GLMM framework. Moreover, because it uses all family samples, FSKAT claims to be more powerful than SKAT that uses only unrelated individuals (founders) in the family data. FSKAT constructs a kinship matrix based on pedigree relationships using the R kinship library. We ran FSKAT correcting for sex and first two PCs.

#### 2.4.4 EPACTS

Efficient and Parallelizable Association Container Toolbox (EPACTS) is a stand-alone software that implements several gene-based statistical tests (CMC, VT and SKAT) and adapts them to complex families by using EMMAX (https://genome.sph.umich.edu/wiki/EPACTS). EPACTS generates a kinship matrix based on BN algorithm and also annotates the genotypic input file and offers filtering tools (frequency and predicted effect of variants) for easier user-selection of variants that go into gene-based analysis. Nonetheless, we used the same set of variants as in other tests, and corrected for sex and first two PCs, to run our analysis with EPACTS.

#### 2.4.5 FarVAT

The Family-based Rare Variant Association Test (FarVAT) (Choi et al. 2014) provides a burden and a variance component test (VT) for extended families, and extends these approaches to the SKAT-O statistic. FarVAT assumes that families are ascertained based on the disease status if family members, and minor allele frequencies between affected and unaffected individuals are compared. FarVAT is implemented in C++ and is computationally efficient. Additionally, if genotype frequencies of affected and unaffected samples are compared to detect the genetic association, it has been shown that the statistical efficiency can be improved by modifying the phenotype; and so FarVAT uses prevalence (Lange and Laird 2002) or Best Linear Unbalanced Predictor (BLUP) (Thornton and McPeek 2007) as covariate to modify the genotype.

#### 2.4.6 PedGene

PedGene (Schaid et al. 2013) is an R package that extends burden and kernel statistics to analyze binary traits in family data, using large-scale genomic data to calculate pedigree relationships. To derive the kernel association statistic and the burden statistic for data that includes related subjects, they take a retrospective view of sampling, with the genotypes considered random.

#### 2.4.7 RVGDT

The Rare Variant Generalized Disequilibrium Test (RVGDT) (He et al. 2017), implemented in Python, differs from the previous methods presented. Instead of using a kernel method to evaluate variants, derives from the generalized disequilibrium test (GDT) which uses genotype differences in all discordant relative pairs to assess associations within a family (Chen et al. Rich 2009). The rare-variant extension of GDT (RVGDT) aggregates a single-variant GDT statistic over a genomic region of interest, which is usually a gene. We ran RVGDT correcting for sex and first two PCs.

#### 2.4.8 RareIBD

RareIBD (Sul et al. 2016) claims to be a program without restrictions on family size, type of trait, whether founders are genotyped, or whether unaffected individuals are genotyped. The method is inspired by non-parametric linkage analysis and looks for a rare variants whose segregation pattern among affected and unaffected individuals is different from the predicted distributions based on Mendelian inheritance and computes a statistic measuring the difference.

## 3 Results

### 3.1 Simulated dataset

Results from the simulated dataset indicate that RVGDT, rareIBD and collapsing-based methods (Burden, CMC and CLP), provided more statistical power than the variance-component methods to detect association of perfectly segregating variants with disease status (**Table 4**).

In an hypothetical scenario of five families in which each one of these families presents perfect segregation with disease status for a different variant within the same gene (5FC×0NFC), transmission-disequilibrium based methods evaluate this association as significant (even after multiple test correction; e.g. RVGDT p-value=0.004; p-value after multiple test correction 0.004×9 = 0.036). RVGDT reaches a ceiling p-value of 1×10^-4^; at 10 families carriers (FC) plus 15 families non-carriers (FNC). RVGDT was unable to produce a p-value smaller than 9×10^-4^, therefore it is not possible to rank or determine the significance of genes with this p-value. Similarly, RareIBD reports the same p-value for all simulated scenarios, which can be an artifact or a flaw of the program. Collapsing-based methods (Burden, CMC and CLP) started with significant p-values for the 5FC×0NFC scenario, but as we added FNC in the analysis, the association became less significant. Then, as we increased the number of FC of segregating variants, the association became more significant. In our analyses, most variance-component tests could not work with the scenarios with only five families carrying the segregating variant; most of the tests only provided p-values once 25 families are included in the analysis (5FC×20FNC). After that, as we increased the number of FC of a segregating variants, the p-value became smaller. SKAT required 15FC×10FNC to report nominally significant p-values, GSKAT required 20FC×5FNC to report statistically significant p-values, FarVAT-CALPHA did not generate significant p-values, except if we used the BLUP correction; FarVAT SKATO reported p-values that were significant at 15FC×10FNC, and at 5FC×20FNC if we used the BLUP correction. P-values from EPACTS-SKAT were not statistically significant after multiple test correction. FSKAT did not deal well with perfectly segregating scenarios; it did not provide p-values for a scenario of only five families all carriers of the segregating variant (5FC×0FNC - FSKAT p-value=NA), and after five families carrying the segregating variant, the program saturated giving no p-value.

Overall, Transmission-disequilibrium tests and collapsing tests were the models that identified these simulated segregating variants as associated with the phenotype; the CMC model provided by FarVAT-BLUP was the one providing most genome-wide significant p-values, even in the 5FC×0FNC scenario.

### 3.2 Candidate genes - APOE

We examined the performance of four gene-sets generated for the *APOE* gene with the twenty-two family-based gene-based methods in our entire familial cohort. Neither the entire set of polymorphic variants (set “gene” in Table 5) nor the set including only rare non-synonymous variants (set “HM” in Table 5) confer risk for these families. The association seems to be driven by the common *APOE* ε2 and ε4 variants, since only when these were considered, either alone (set “ε2ε4” in Table 5) or in conjunction with the rest of rare non-synonymous variants (set “HM-ε2ε4” in Table 5), most of the tests yielded a significant p-value (after multiple test correction). Only EPACTS-SKAT did not consider the *APOE* ε2 and ε4 variants as significantly associated, after multiple test correction, with our dataset (**Table 5**). The most significant association for *APOE* ε2 and ε4 variants was reported by FarVAT-CMC test.

**Table 5.**
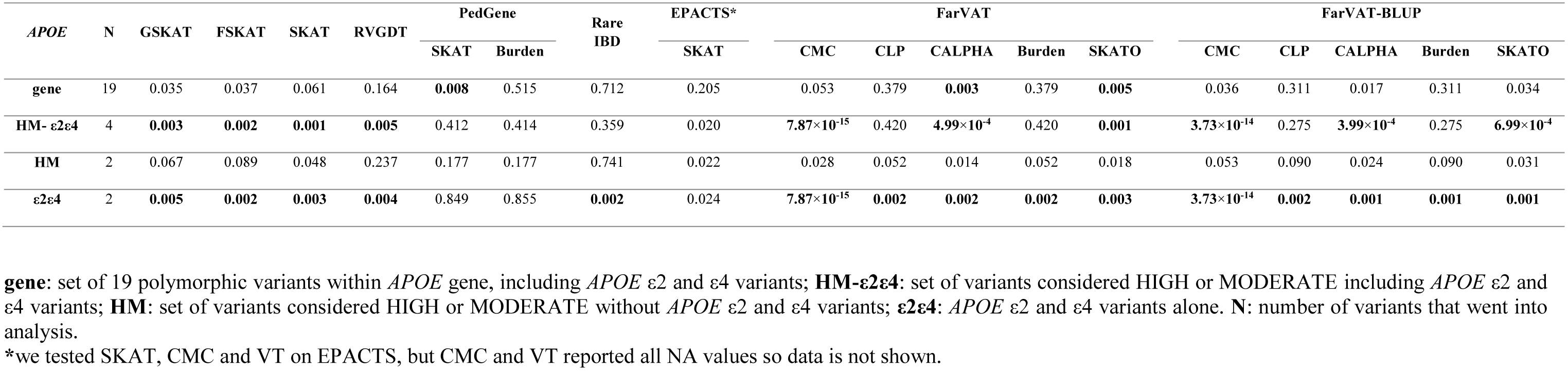
Gene-based p-values for the *APOE* gene under different gene-set scenarios for the gene-based methods tested in the entire dataset (N=1235, 285 families). In the analysis, only nonsynonymous variants (only SNVs) with a MAF<0.01, and the APOE ε2 and ε4, were considered and we adjusted by sex and PCAs. Highlighted in bold, significant p-values after multiple test correction.

### 3.3 Genome-wide analyses

Overall, we examined eight software and over 22 algorithms for genome-wide association analysis in our extended family dataset of 285 families and 1235 non-hispanic white individuals. We only included in the analysis non-synonymous SNPs with a MAF ≤ 1% and we corrected per sex and first two PCs. All 22 algorithms were run using the same input dataset. The results for these 22 algorithms are described grouped per category, as detailed in the following sections. First, we compared the correction effect provided by four kinship matrices (**Figure 3A**). Second, we compare the performance of nine variance-component software and algorithms (**Figure 3B**). Third is the comparison of eight collapsing software and algorithms. Fourth, we compare two transmission-disequilibrium tests. We conclude the results section by providing a summary of the pros and cons encountered while running these methods. Overall, most of the gene-based methods tested seemed quite deflated. Only PedGene, FarVAT and Rare-IBD seem to provide values closer or above the expected under the null hypothesis. The most efficient in terms of power and p-value inflation appears to be FarVAT with BLUP correction.

**Figure 3.**
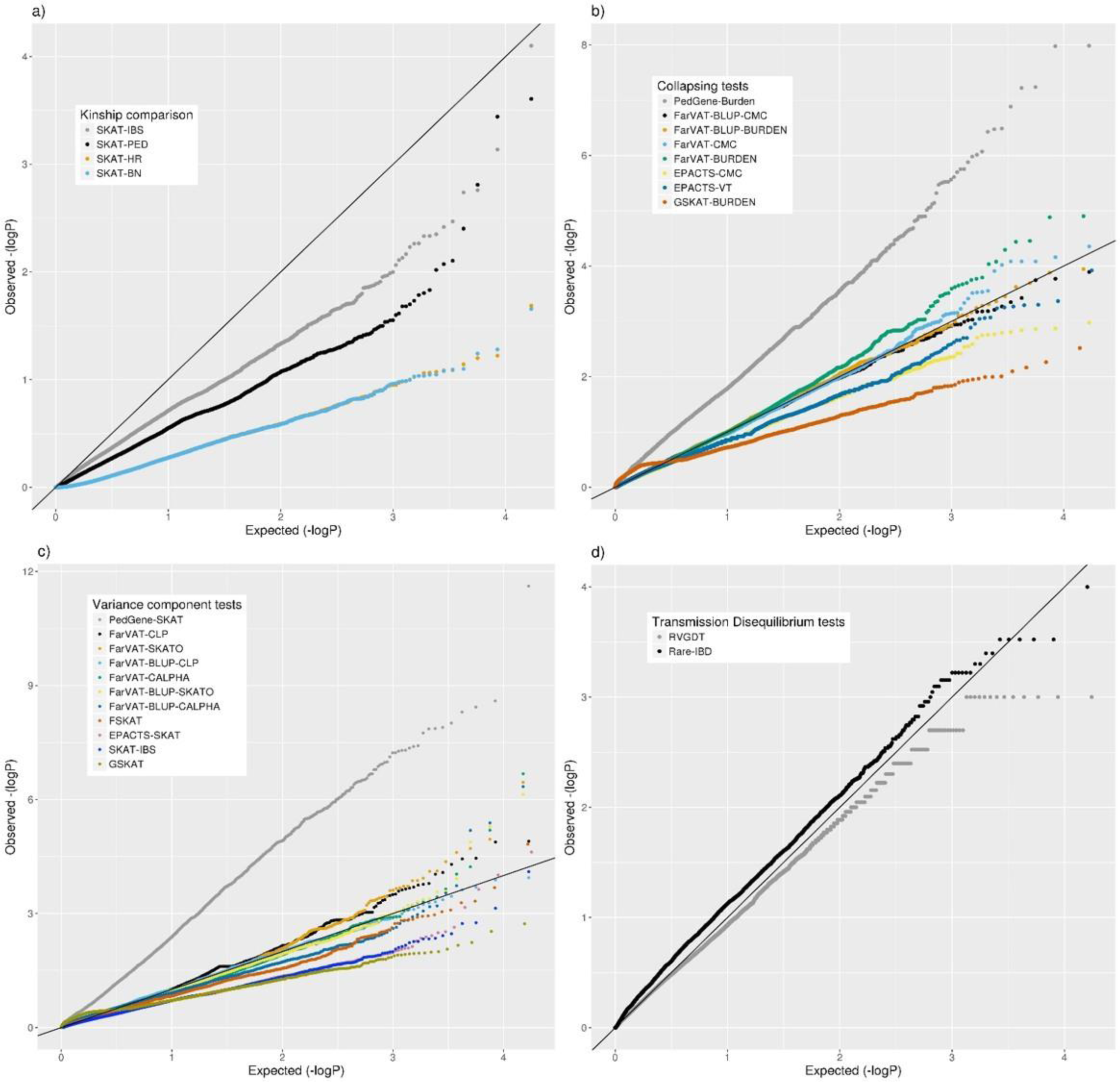
Quantile-quantile (QQ) plots from different family-based gene-based methods for all nonsynonymous variants with a MAF <1% in our family-based dataset. a) Comparison of SKAT test using different kinship matrices: pedigree calculation (PED), Identity By Similarity (IBS) estimation, Balding-Nichols (BN) estimation, and the kinship generated by EPACTS (HR). c) Comparison of different collapsing tests: GSKAT, EPACTS, FarVAT and PedGene. b) Comparison of different variance-component gene-based methods: GSKAT, FSKAT, SKAT, EPACTS, FarVAT and PedGene. d) Comparison of transmission disequilibrium tests: RVGDT and RareIBD.

#### 3.3.1 Kinship matrices

We tested the correction provided by four kinship matrices using the SKAT method with EMMAX correction implemented in the R package SKATv2. The four kinship matrices tested were pedigree calculation (PED), Identity By State (IBS) estimation, Balding-Nichols (BN) estimation, and the kinship generated by EPACTS (HR) which is also based on BN algorithm (**Figure 3A**). **Table S3** offers a comparison of these kinships for FAM#1 and FAM#2 of our simulated dataset. For these analyses, we ran the SKAT-EMMAX method in our entire dataset, gene-wide and calculated a QQ plot and inflation factor (λ) to obtain a general ideal of the behavior of each matrix. Matrices based on the BN algorithm seemed to have a similar performance (SKAT-BN λ=0.038, SKAT-HR λ=0.039, **Table 6**) although their concordance was lower than expected given they are based on the same algorithm (Pearson correlation (Pc)=0.85; Spearman correlation (Sc)=1). Although the PED matrix generates a more restrictive correction than the IBS matrix (SKAT-PED λ = 0.36, SKAT-IBS λ=0.67, **Table 6**), these two tests have a similar overall performance as the p-values for the different genes are highly correlated (Pc=0.97; Sc=0.98), making the PED matrix a good surrogate for the IBS matrix. Finally, there were clear performance differences between the BN-type matrices (BN and HR) and the IBS-type matrices (IBS and PED), exemplified by the different top candidate genes (*NR1D1* for BN-type matrices and *CHRD* for IBS-type matrices) and by the correlation algorithms (SAKT-IBS vs SKAT-BN Pc=0.8; Sc=0.89). Overall, we found that the IBS matrix provided to our dataset the best balance between covariance-correction and overcorrection.

**Table 6.**
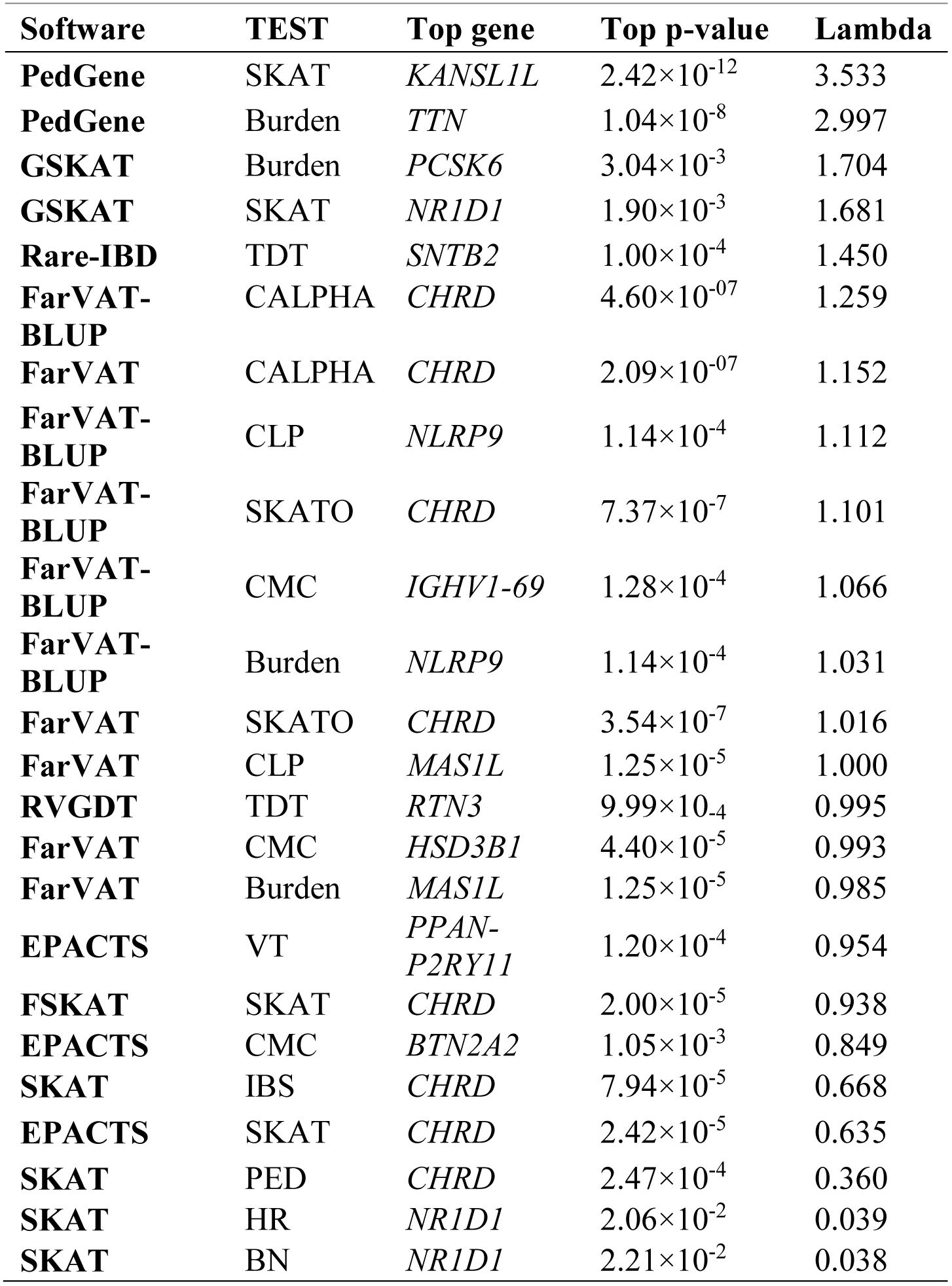
Top results for all gene-based methods tested. Top gene, p-value and lambda for each test is given, ordered by lambda value.

#### 3.3.2 Collapsing tests

The collapsing methods tested from four different software (PedGene, FarVAT, EPACTS and GSKAT) were Burden, CMC and VT (**Figure 3c**). In order to compare the different tests, we followed a similar approach as above, and we ran the different software with the same imputed file and compared the λ.

In our analyses, the burden test by GSKAT presented the most deflated values; although the lambda does not illustrate so (GSKAT-Burden λ= 1.71, **Table 6**) because of the initial inflation among the low or non-significant genes. EPACTS-CMC (λ= 0.85) and EPACTS-VT (λ=0.95) provided values closer to the expected, and despite their QQ-plots seem to follow a similar trend, their correlation is weak (Pc=0.54; Sc=0.68), pointing to different top genes. The Burden and CMC methods by FarVAT and FarVAT-BLUP provided p-values closest to the expected (FarVAT-Burden λ=0.98; FarVAT-CMC λ=0.99, FarVAT-BLUP-Burden λ=1.03; FarVAT-BLUP-CMC sλ=1.07). The correlation for the gene p-values was higher between results generated by the same method (FarVAT-BLUP-CMC vs FarVAT-BLUP-Burden Pc=0.99; Sc=0.96; FarVAT-CMC vs FarVAT-Burden Pc=0.98; Sc=0.97) than between results generated using the same algorithm (FarVAT-BLUP-CMC vs FarVAT-CMC Pc=0.88; Sc=0.8; FarVAT-BLUP-Burden vs FarVAT-Burden Pc=0.85; Sc=0.77). PedGene in the burden model is the software that provided most significant p-values; however, these are clearly inflated compared to the predicted p-values (Pedgene-Burden λ=2.99, **Table 6**) and its results were not correlated with any other Collapsing test (Pc and Sc values < 0.1).

#### 3.3.3 Variance component tests

This subset included all the Variance component-based methods available, CLP, CALPHA and SKAT, from six different software: PedGene, FarVAT, FSKAT, EPACTS, SKAT and GSKAT (**Figure 3c**). GSKAT was the software presenting more deflated values though the lambda does not illustrate this (GSKAT-SKAT λ = 1.681, **Table 6**) because of the initial inflation among the low or non-significant genes. GSKAT was followed by SKAT and EPACTS which showed similar λ and performance-values for each gene (Pc=0.8, Sc=0.8, **Figure 4**). The CLP, CALPHA and SKATO methods by FarVAT and FarVAT-BLUP provided p-values closest to the expected (FarVAT-CLP λ=1.00; FarVAT-CALPHA λ =1.15; FarVAT-SKATO λ=1.02, FarVAT-BLUP-CLP λ=1.11; FarVAT-BLUP-CALPHA λ=1.26; FarVAT-BLUP-SKATO λ=1.10). FarVAT-CALPHA, FarVAT-SKATO, FarVAT-BLUP-CALPHA and FarVAT-BLUP-SKATO pointed to the same top candidate gene (*CHRD*) (Table 6), although the overall p-value correlation is lower than expected considering they are based on the same algorithm (FarVAT-SKATO vs FarVAT-BLUP-SKATO Pc=0.6, Sc=0.7; FarVAT-CALPHA vs FarVAT-BLUP-CALPHA Pc=0.82 Sc=0.82, **Figure 4**). On the other hand, and despite the fact that FarVAT-CLP and FarVAT-BLUP-CLP have higher correlation (Pc=0.85, Sc=0.77), these two tests point to different top genes (FarVAT-CLP top gene is *MAS1L*, and FarVAT-BLIP-CLP top gene is *NLRP9*). PedGene in the SKAT model is the software that provided the most significant p-values, but we can observe how these are inflated (Pedgene-SKAT λ=3.53, **Table 6**) and that its correlation with other variance component tests is low to null (Pc and Sc values < 0.2).

**Figure 4.**
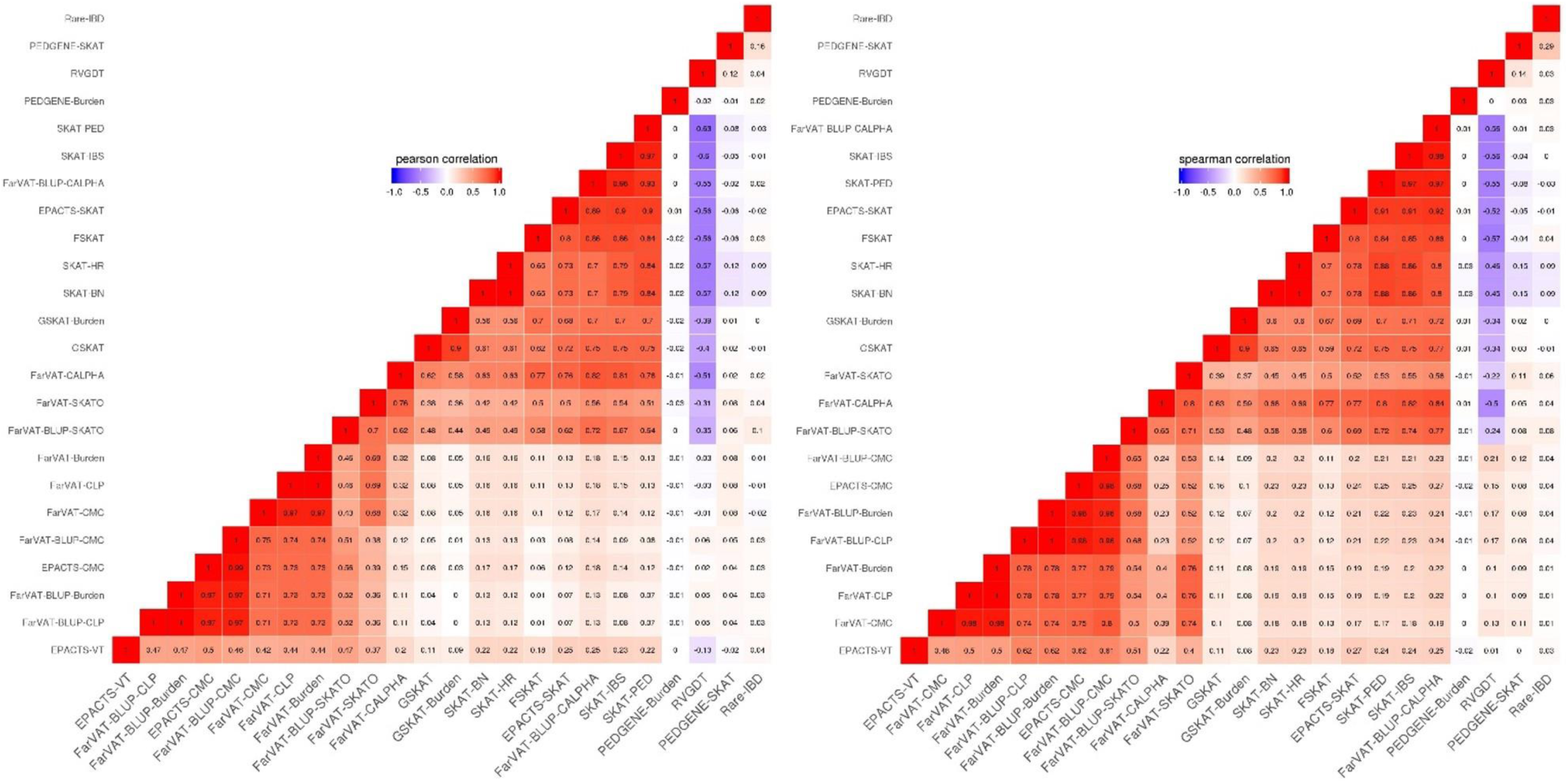
Correlation plots from different family-based gene-based methods for genes with a p-value ≤ 0.005. a) Pearson correlation correlates genes according to their p-values. b) Spearman correlation correlates genes according to their rankings.

#### 3.3.4 Transmission disequilibrium tests

We have tested two transmission disequilibrium tests, RVGDT and Rare-IBD, which are designed to account for large extended families of arbitrary structure (**Figure 3d**). Of these two, RVGDT is the test that more closely approached the expected under the null (λ=0.99), whereas Rare-IBD provided slightly inflated p-values (λ=1.450, **Table 6**). The correlation between these two methods was very low (Pearson correlation = 0.23, Spearman correlation = 0.17). A common issue with both methods is that we could see some stratification towards more significant p-values which made it difficult to determine a top significant gene.

#### 3.3.5 PROS and COSN of the different gene-based methods

Among all the methods tested, EPACTS and FarVAT are the most user-friendly, time-efficient and versatile software. EPACTS is an all-in-one package that annotates the input file, generates the kinship matrix and performs gene-based analysis under different conditions (minor allele frequency and predicted functionality of the variant) with only tag specification. In addition, the program can be run on a genome-wide base or at smaller scale given genes or regions specified by the user. FarVAT can generate the kinship matrix by either using the pedigree relationships or using the genetic relationship among individuals. It does not annotate the input file and requires that the user provide their own set of genes and variants per gene to analyze; it allows the user to choose between BLUP (best linear unbiased prediction) or prevalence to estimate and incorporate random effects on the phenotype. FarVAT has initial conditioning that only takes founder-based MAF, i.e. when a genetic variant has its minor alleles only in non-founders (offspring), these numbers will not be counted. This is a big difference with respect to the other programs that take into account all variants regardless of their presence in founders or not. Since for many of our families we only had genetic data for siblings, i.e. we did not have genetic data for founders, we ran FarVAT with the “–freq all” option, so all variants would be included regardless if they are present in founders or not.

FSKAT, GSKAT and SKAT require of some R knowledge from the user, and are less flexible. For FSKAT and GSKAT the user has to provide a genotype, a phenotype, and a gene-set file. For SKAT the user has to additionally provide the kinship matrix. Because these programs were designed to run on a per gene basis, these take longer to compute and to be run on a genome-wide level than EPACTS or FarVAT, even if the user parallelizes computation. PedGene is also an R package that requires a genotype, a phenotype file with complete pedigree information (to generate the kinship matrix), and a gene-set file. PedGene provides phenotype adjustment by logistic regression on the trait of interest, but it does not allow for extra covariates, which prohibits correction by multiple PCs or other variables. RVGDT is a python based program, quite user-friendly since it is operated with simple command-line but is limited in its options. Similar to FSKAT, GSKAT and SKAT, it is designed to be run on a per-gene basis for which loops and parallelization have to be set up for genome-wide testing. The same goes for RareIBD which requires a genotype, a phenotype, and a Kinship coefficient file for each gene that the user wants to test. For each gene the program computes first statistics for each founder within each family and then calculates the gene-based p-value. The first step of this process can easily take between three to five minutes for families with less than 100 individuals; hence, the overall time for one gene is directly dependent on the number of families to test and the time required for a genome-wide analysis is proportional to the number of genes being tested. Although it is possible to parallelize the jobs using a high-performance cluster (if available) this program is the slowest of all tested.

One of the major drawbacks we found is that some of these programs do not accept missing data (FSKAT or RareIBD) or will not generate a p-value if the gene set contains only one variant (GSKAT, PedGene or FarVAT). FSKAT does not accept missing data, and although it calculates p-values for genes that only have one informative SNP (2154 one-SNP-gene), there were at least 75 (3.26%) of these one SNP-genes for which the returned p-value was “2”. GSKAT did not provide p-values for more than 1,875 one-SNP-genes. Pedgene also had trouble generating p-values for 44 one-SNP-genes out of a total of 1,916 singletons. FarVAT did not generate a p-value for the 1,875 one-SNP-genes using the Burden and SKATO models but it generated p-values using the CMC and CLP models for the same 1,875 one-SNP-genes.

### 3.4 Candidate genes for FASe project

Our results indicate that transmission disequilibrium tests identify genes that have a Mendelian behavior, whereas collapsing and variance-component tests identify genes that confer risk for disease. Therefore, we decided to combine and compare results from all approaches to identify the genes with most consistent results (**Table 7**).

**Table 7.**
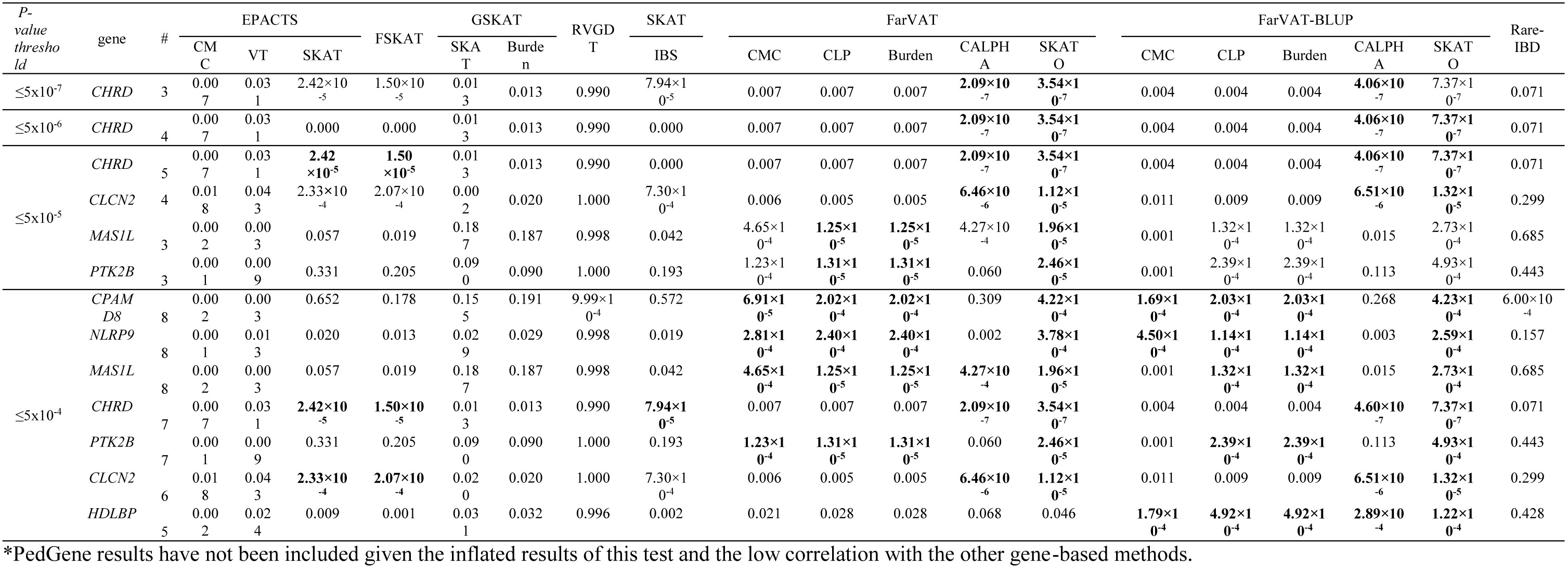
Most frequent genes, within p-value threshold category, across the different gene-based family-based methods tested. Highlighted in bold the tests with significant p-value according to threshold category.

PEDGENE provided the most significant p-values for *NTN5* (Pedgene-Burden p-value = 5.80×10^-8^; Pedgene-SKAT p-value = 1.26×10^-8^) and *ANKRD42* (Pedgene-Burden p-value = 3.62×10^-7^; Pedgene-SKAT p-value = 1.16×10^-7^). However, the inflated p-values observed and low correlation with any of the other software tested using the same algorithms makes us suspicious of the validity of these results.

*CHRD* was the gene with the third most significant p-value. *CHRD* had a p-value ≤5×10^-7^ in three different models (FarVAT-CALPHA, FarVAT-SKATO, FarVAT-BLUP-CALPHA). In addition, as we lowered the considered p-value threshold we found that more tests identified *CHRD* as a potential candidate gene associated with AD. When we lowered the threshold to suggestive genome-wide p-value (p-value≤5×10^-4^) we found that seven different models identified *CHRD* as a gene significantly associated with AD. Following the same method we found that *CLCN2, MAS1L* and *PTK2B* had p-values ≤ 5×10^-05^ in at least three tests, and if we lowered the threshold to ≤5×10^-4^ p-value, these genes were identified as significant by at least three additional tests.

Among genes with a p-value ≤ 5×10^-04^; *CPAMD8* was identified by at least nine gene-based methods (FarVAT, FarVAT-BLUP and PedGene). The exact p-value for *CPAMD8* could not be estimated by RVGDT as it showed a p-value of 9×10^-04^, which is the most significant p-value provided by this test. Therefore, we cannot conclude that *CPAMD8* presented a p-value ≤ 5×10^-04^ by RVGDT. *CHRD*, *CLCN2, MAS1L*, *PTK2B* and *CPAMD8*, *NLRP9*, and *HDLBP* were also potential novel candidate genes for familial LOAD as they had p-values ≤ 5×10^-04^ using at least five or more tests (**Table 7**).

Since these were identified by multiple gene-based methods, we wanted to determine whether any of these seven candidate genes are involved in known AD pathways. Common variants in *PTK2B* have been associated with AD risk at genome-wide level (J.-C. Lambert et al. 2013). Our results indicate there are additional low-frequency and rare non-synonymous variants in *PTK2B* that are associated with AD risk in late-onset families.

We used the GeneMANIA (http://pages.genemania.org/) algorithm on the seven candidate genes (*CHRD, MAS1L, PTK2B, CPAMD8, NLRP9, CLCN2* and *HDLBP*) along with known AD-related genes (APP, *PSEN1, PSEN2, APOE, TREM2, PLD3, ADAM10*) which represent some of the AD genes and pathways (APP-metabolism and immune response). GeneMANIA is a software that looks for relationships among a list of given genes by searching within multiple publicly available biological datasets. These datasets include protein-protein, protein-DNA and genetic interactions, pathways, reactions, gene and protein expression data, protein domains and phenotypic screening profiles. We found that our candidate genes have genetic interactions and co-localization with known AD genes. *CHRD* and *PTK2B* are involved in “regulation of cell adhesion” like *ADAM10; PTK2B* is involved in “regulation of neurogenesis” like *APOE* and “perinuclear region of cytoplasm” like *APP, PSEN1* and *PSEN2*. Finally, *CLCN2* and *PTK2B* are connected through “regulation of ion transport” (Figure 5).

**Figure 5.**
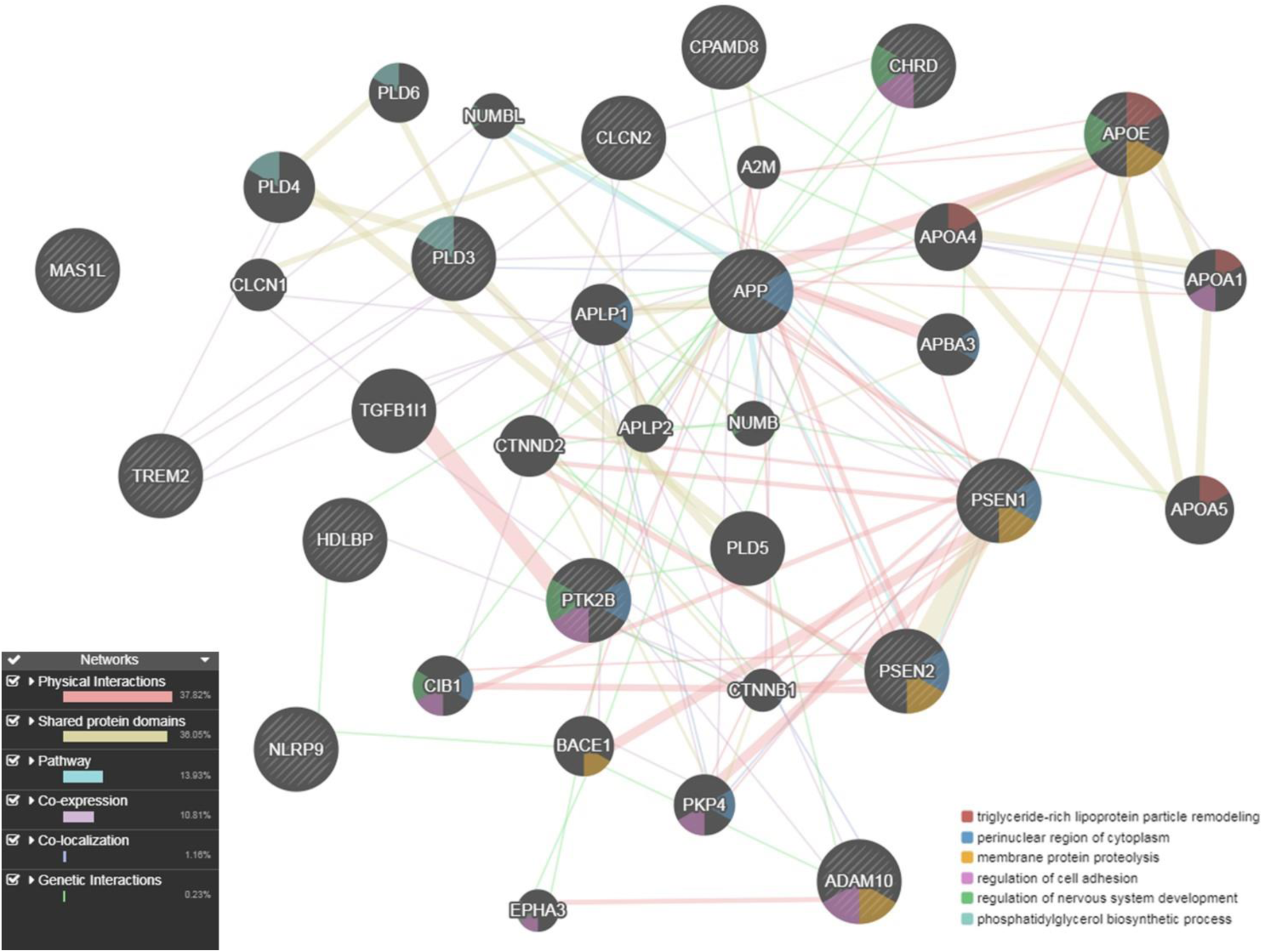
Gene network for the seven candidate genes (*CHRD, CLCN2, CPAMD8, HDLBP, MAS1L, NLRP9* and *PTK2B*) with multiple evidence of a p-value ≤ 5×10^-04^, anchored with known AD genes (*APP, PSEN1, PSEN2, APOE, TREM2, ADAM10, PLD3*), as described by GeneMania.

## 4 Discussion

The remaining missing heritability in AD, and in many complex diseases, may be found in very rare-variants for which discovery will require either large datasets (eg. the ADSP Discovery Phase which has over 10,000 sequenced individuals) or datasets enriched for rare variants (such as families with history of AD). In this study, we present the most comprehensive performance analyses for multiple gene-based methods in 285 families with AD. Some of the current methods available are underpowered or too restrictive to detect genes significantly associated with this disease (**Figure 4**). Results from our simulated data (**Table 4**) show that only certain highly restricted scenarios provide gene-wide significant p-values in a family-based analysis; whereas, similar scenarios in a case-control study would result in gene-wide p-values. To circumvent this power issue, we relied on the combination of multiple evidence towards the same gene.

One key aspect to adapt gene-based analyses to a family-based context is to account for the population stratification and hidden relatedness that may appear due to the inherent nature of the dataset. To take into account this issue, gene-based algorithms must incorporate kinship matrices to model the relationships among samples. Therefore, an appropriate estimate of the kinship matrix is of utmost importance. In this work we show how different relationship matrices influence results. We tested the three most common types of kinship matrix, pedigree reconstruction (PED), identity by state (IBS), and Balding-Nichols (BN). We show that for a situation of complex incomplete families, correction using PED or BN matrices will lead to an overcorrection of the relationships decreasing the power of these tests (**Table 6**, **Figure 4A**).

In order to choose the best gene-based algorithm for analysis, it is important to take into account the nature (impact and directionality) of the variants that are being included in the test. Collapsing tests are powerful when a large proportion of variants are causal and effects are in the same direction. Variance-component tests are supposed to be more powerful than collapsing tests because these allow for admixture of risk and protective variants within the region being tested (Ionita-Laza et al. 2013). It is not practical to account for the nature of the variants included in each gene-set, and the true disease model is unknown and variable; hence, omnibus or combined tests such as SKAT-O would be desirable for genome-wide studies (Lee et al. 2012); however, most family-based methods do not incorporate the SKAT-O algorithm, except for FarVAT. Therefore, the best approach to perform genome-wide rare variant discovery is to combine different algorithms and look for common signatures across the tests performed. Nonetheless, we are aware that running all available tests is a time-consuming task that requires additional expertise and resources. In our analyses FarVAT, with the BLUP adjustment, provide the best results in terms of significant p-values and inflation, for genome-wide gene-based analysis; it is a fast software that provides results from multiple tests at the same time. The R version of SKAT or EPACTS, would be alternative valid choices, taking into account that these overcorrect and the p-value threshold should be lowered.

In this study, we identified *CHRD* as a candidate gene with a genome-wide significant p-value (5× 10^-07^) reported by three tests, and another six genes that had a suggestive genome-wide p-value < 5×10^-04^ in at least five and up to nine of the different test performed: *CLCN2, CPAMD8, HDLBP, MAS1L, NLRP9* and *PTK2B*. In addition, these genes seem to have direct and indirect interactions (genetic interaction, co-localization or shared function) with known AD genes (APP, *PSEN1, PSEN2, APOE, TREM2, PLD3* and *ADAM10*).

*CHRD*, chordin, is a developmental protein, highly conserved, inhibiting the ventralizing activity of bone morphogenetic proteins, active during gastrulation, expressed in fetal and adult liver and cerebellum, associated with Cornelia de Lange syndrome (Smith et al. 1999). *CLCN2*, chloride voltage-gated channel 2, has several functions including the regulation of cell volume; membrane potential stabilization, signal transduction and transepithelial transport. It has been associated with different epilepsy modes (Saint-Martin et al. 2009; Cukier et al. 2014) and leukoencephalopathy (Gaitán-Peñas et al. 2017). *CHRD* and *CLCN2* show co-expression which could be due to their close location, both belong to a gene cluster at 3q27. Interestingly, *CLCN2* shows co-expression with *TREM2*, which other than being a risk gene for AD, is known to cause leukoencephalopathy in the PLOSL (polycystic lipomembranous osteodysplasia with sclerosing leukoencephalopathy) form, also known as Nasu-Hakola disease.

*PTK2B*, was described as a GWAs hit locus in the largest GWAs meta-analysis conducted to date (Lambert et al. 2013), and later corroborated by others (Wang et al. 2015; Beecham et al. 2014). The protein encoded by *PTK2B* is a member of the focal adhesion kinase (FAK) family that can be activated by changes in intracellular calcium levels, which are disrupted in AD brains. Its activation regulates neuronal activity such as mitogen-activated protein kinase (MAPK) signaling (Rosenthal and Kamboh 2014). *PTK2B* could also be involved in hippocampal synaptic function (Lambert et al. 2013). Although there is no co-expression or genetic interaction between *CLCN2* and *PTK2B*, both are involved in regulation of ion transport. Additionally, *PTK2B* is involved in regulation of lipidic metabolic processes, like APOE, a cholesterol-related gene. Despite no association has yet been reported between *APOE* and *HDLBP*, the High-Density Lipoprotein Binding Protein plays a role in cell sterol metabolism, protecting cells from over-accumulation of cholesterol, which has been reported as risk factor for atherosclerotic vascular diseases.

*CPAMD8* causes a Unique Form of Autosomal-Recessive Anterior Segment Dysgenesis (Cheong et al. 2016). No shared pathway association was found between *CPAMD8* and the known AD genes, but it seems to have a genetic interaction with *APP* (Lin et al. 2010). In our study *CPAMD8* was identified as a candidate gene (with p-value < 1×10^-4^) for AD by at least nine gene-based methods from different software, and we found that several variants within this gene show varying degrees of perfect segregation in more than twenty families. Variant p.(Ser1103Ala) segregates with disease status in two families with two and three carriers respectively, and is present in another two families. Variant p.(His465Arg) segregates with disease status in five families with two or three carriers per family and is present in another 11 families. Variant p.(Arg1380Cys) is private to a family with three carriers, p.(Ala1492Pro) is private to a family with five carriers, and p.(Val521Met) is private to a family with three carriers.

We have reviewed over 22 algorithms from eight different software available for the gene-based analysis in complex families. After a thorough examination of these tests performance under different scenarios, we present a methodology to identify genes associated with the studied phenotype. We have applied this methodology to 285 European-American families affected with late onset Alzheimer disease (LOAD). We have identified six candidate genes with suggestive or significant genome-wide p-values and we are confident that some of these genes are truly involved on AD pathology.

## 6 Author contributions statement

## 8 Acknowledgments

We thank contributors who collected samples used in this study, as well as patients and their families, whose help and participation made this work possible. Members of the National Institute on Aging Late-Onset Alzheimer Disease/National Cell Repository for Alzheimer Disease (NIA-LOAD NCRAD) Family Study Group include the following: Richard Mayeux, MD, MSc; Martin Farlow, MD; Tatiana Foroud, PhD; Kelley Faber, MS; Bradley F. Boeve, MD; Neill R. Graff-Radford, MD; David A. Bennett, MD; Robert A. Sweet, MD; Roger Rosenberg, MD; Thomas D. Bird, MD; Carlos Cruchaga, PhD; and Jeremy M. Silverman, PhD.

The Alzheimer’s Disease Sequencing Project (ADSP) is comprised of two Alzheimer’s Disease (AD) genetics consortia and three National Human Genome Research Institute (NHGRI) funded Large Scale Sequencing and Analysis Centers (LSAC). The two AD genetics consortia are the Alzheimer’s Disease Genetics Consortium (ADGC) funded by NIA (U01 AG032984), and the Cohorts for Heart and Aging Research in Genomic Epidemiology (CHARGE) funded by NIA (R01 AG033193), the National Heart, Lung, and Blood Institute (NHLBI), other National Institute of Health (NIH) institutes and other foreign governmental and non-governmental organizations. The Discovery Phase analysis of sequence data is supported through UF1AG047133 (to Drs. Schellenberg, Farrer, Pericak-Vance, Mayeux, and Haines); U01AG049505 to Dr. Seshadri; U01AG049506 to Dr. Boerwinkle; U01AG049507 to Dr. Wijsman; and U01AG049508 to Dr. Goate and the Discovery Extension Phase analysis is supported through U01AG052411 to Dr. Goate, U01AG052410 to Dr. Pericak-Vance and U01 AG052409 to Drs. Seshadri and Fornage. Data generation and harmonization in the Follow-up Phases is supported by U54AG052427 (to Drs. Schellenberg and Wang).

The ADGC cohorts include: Adult Changes in Thought (ACT), the Alzheimer’s Disease Centers (ADC), the Chicago Health and Aging Project (CHAP), the Memory and Aging Project (MAP), Mayo Clinic (MAYO), Mayo Parkinson’s Disease controls, University of Miami, the Multi-Institutional Research in Alzheimer’s Genetic Epidemiology Study (MIRAGE), the National Cell Repository for Alzheimer’s Disease (NCRAD), the National Institute on Aging Late Onset Alzheimer's Disease Family Study (NIA-LOAD), the Religious Orders Study (ROS), the Texas Alzheimer’s Research and Care Consortium (TARC), Vanderbilt University/Case Western Reserve University (VAN/CWRU), the Washington Heights-Inwood Columbia Aging Project (WHICAP) and the Washington University Sequencing Project (WUSP), the Columbia University Hispanic-Estudio Familiar de Influencia Genetica de Alzheimer (EFIGA), the University of Toronto (UT), and Genetic Differences (GD).

The CHARGE cohorts are supported in part by National Heart, Lung, and Blood Institute (NHLBI) infrastructure grant HL105756 (Psaty), RC2HL102419 (Boerwinkle) and the neurology working group is supported by the National Institute on Aging (NIA) R01 grant AG033193. The CHARGE cohorts participating in the ADSP include the following: Austrian Stroke Prevention Study (ASPS), ASPS-Family study, and the Prospective Dementia Registry-Austria (ASPS/PRODEM-Aus), the Atherosclerosis Risk in Communities (ARIC) Study, the Cardiovascular Health Study (CHS), the Erasmus Rucphen Family Study (ERF), the Framingham Heart Study (FHS), and the Rotterdam Study (RS). ASPS is funded by the Austrian Science Fond (FWF) grant number P20545-P05 and P13180 and the Medical University of Graz. The ASPS-Fam is funded by the Austrian Science Fund (FWF) project I904),the EU Joint Programme - Neurodegenerative Disease Research (JPND) in frame of the BRIDGET project (Austria, Ministry of Science) and the Medical University of Graz and the Steiermärkische Krankenanstalten Gesellschaft. PRODEM-Austria is supported by the Austrian Research Promotion agency (FFG) (Project No. 827462) and by the Austrian National Bank (Anniversary Fund, project 15435. ARIC research is carried out as a collaborative study supported by NHLBI contracts (HHSN268201100005C, HHSN268201100006C, HHSN268201100007C, HHSN268201100008C, HHSN268201100009C, HHSN268201100010C, HHSN268201100011C, and HHSN268201100012C). Neurocognitive data in ARIC is collected by U01 2U01HL096812, 2U01HL096814, 2U01HL096899, 2U01HL096902, 2U01HL096917 from the NIH (NHLBI, NINDS, NIA and NIDCD), and with previous brain MRI examinations funded by R01-HL70825 from the NHLBI. CHS research was supported by contracts HHSN268201200036C, HHSN268200800007C, N01HC55222, N01HC85079, N01HC85080, N01HC85081, N01HC85082, N01HC85083, N01HC85086, and grants U01HL080295 and U01HL130114 from the NHLBI with additional contribution from the National Institute of Neurological Disorders and Stroke (NINDS). Additional support was provided by R01AG023629, R01AG15928, and R01AG20098 from the NIA. FHS research is supported by NHLBI contracts N01-HC-25195 and HHSN268201500001I. This study was also supported by additional grants from the NIA (R01s AG054076, AG049607 and AG033040 and NINDS (R01 NS017950). The ERF study as a part of EUROSPAN (European Special Populations Research Network) was supported by European Commission FP6 STRP grant number 018947 (LSHG-CT-2006-01947) and also received funding from the European Community's Seventh Framework Programme (FP7/2007-2013)/grant agreement HEALTH-F4-2007-201413 by the European Commission under the programme “Quality of Life and Management of the Living Resources” of 5th Framework Programme (no. QLG2-CT-2002-01254). High-throughput analysis of the ERF data was supported by a joint grant from the Netherlands Organization for Scientific Research and the Russian Foundation for Basic Research (NWO-RFBR 047.017.043). The Rotterdam Study is funded by Erasmus Medical Center and Erasmus University, Rotterdam, the Netherlands Organization for Health Research and Development (ZonMw), the Research Institute for Diseases in the Elderly (RIDE), the Ministry of Education, Culture and Science, the Ministry for Health, Welfare and Sports, the European Commission (DG XII), and the municipality of Rotterdam. Genetic data sets are also supported by the Netherlands Organization of Scientific Research NWO Investments (175.010.2005.011, 911-03-012), the Genetic Laboratory of the Department of Internal Medicine, Erasmus MC, the Research Institute for Diseases in the Elderly (014-93-015; RIDE2), and the Netherlands Genomics Initiative (NGI)/Netherlands Organization for Scientific Research (NWO) Netherlands Consortium for Healthy Aging (NCHA), project 050-060-810. All studies are grateful to their participants, faculty and staff. The content of these manuscripts is solely the responsibility of the authors and does not necessarily represent the official views of the National Institutes of Health or the U.S. Department of Health and Human Services.

The three LSACs are: the Human Genome Sequencing Center at the Baylor College of Medicine (U54 HG003273), the Broad Institute Genome Center (U54HG003067), and the Washington University Genome Institute (U54HG003079).

Biological samples and associated phenotypic data used in primary data analyses were stored at Study Investigators institutions, and at the National Cell Repository for Alzheimer’s Disease (NCRAD, U24AG021886) at Indiana University funded by NIA. Associated Phenotypic Data used in primary and secondary data analyses were provided by Study Investigators, the NIA funded Alzheimer’s Disease Centers (ADCs), and the National Alzheimer’s Coordinating Center (NACC, U01AG016976) and the National Institute on Aging Genetics of Alzheimer’s Disease Data Storage Site (NIAGADS, U24AG041689) at the University of Pennsylvania, funded by NIA, and at the Database for Genotypes and Phenotypes (dbGaP) funded by NIH. This research was supported in part by the Intramural Research Program of the National Institutes of health, National Library of Medicine. Contributors to the Genetic Analysis Data included Study Investigators on projects that were individually funded by NIA, and other NIH institutes, and by private U.S. organizations, or foreign governmental or nongovernmental organizations.

